# The association between oscillatory burst features and human working memory accuracy

**DOI:** 10.1101/2024.12.06.622989

**Authors:** Brian C. Kavanaugh, Megan M. Vigne, Ryan Thorpe, Christopher Legere, W. Luke Acuff, Noah Vaughan, Eric Tirrell, Saskia Haegens, Linda L. Carpenter, Stephanie R. Jones

## Abstract

Oscillatory power across multiple frequency bands has been associated with distinct working memory (WM) processes. Recent research has shown that previous observations based on averaged power are driven by the presence of transient, oscillatory burst-like events, particularly within the alpha, beta, and gamma bands. However, the interplay between different burst events in human WM is not well understood. The current EEG study aimed to investigate the dynamics between alpha (8-12 Hz)/beta (15-29 Hz) and high frequency activity (HFA; 55-80 Hz) bursts in human WM, particularly burst features and error-related deviations during the encoding and maintenance of WM in healthy adults. Oscillatory burst features within the alpha, beta and HFA bands were examined at frontal and parietal electrodes in healthy young adults during a Sternberg working memory task. Averaged power dynamics were driven by oscillatory burst features, most consistently the burst rate and burst power. Alpha/beta and HFA bursts displayed complementary roles in WM processes, in that alpha and beta bursting decreased during encoding and increased during delay, while HFA bursting had the opposite pattern, i.e., increased during encoding and decreased during the delay. Critically, weaker variation in burst dynamics across stages was associated with incorrect responses and impaired overall task performance. Together, these results indicate that successful human WM is dependent on the rise and fall interplay between alpha/beta and HFA bursts, with such burst dynamics reflecting a novel target for the development of treatment in clinical populations with WM deficits.

## 1. INTRODUCTION

Working memory (WM) is a foundational component of executive function that reflects the process of holding information ‘in mind’ to execute goal-directed behaviors (Diamond, 2013). Based on classic lesion studies, WM was originally thought to be solely centralized in the dorsolateral prefrontal cortex (dlPFC) (Lara & Wallis, 2015). It is now known that the posterior parietal cortex (PPC) also plays a distinct role in WM within the broader central executive network (i.e., frontoparietal network) (Niendam et al., 2012). Recent research has found that the PPC is responsible for encoding the spatial or sensory aspects of stimuli, while the PFC is responsible for executing cognitive control demands (e.g., categorization, filtering). Together, this reflects a parietal-to-frontal feedforward mechanism of spatial signaling as well as a frontal-to-parietal feedback mechanism of control signaling (Crowe et al., 2013; Goodwin et al., 2012; Murray et al., 2017). Furthermore, there is evidence that the PPC is not only involved in aspects of encoding sensory information, but also directly involved in control-related demands of WM (Esterman et al., 2009; Goodwin et al., 2012; Koenigs et al., 2009).

Numerous prior studies have investigated spectral activity underlying WM using scalp electroencephalography (EEG) or magnetoencephalography (MEG) in humans, and most traditional frequency bands have been implicated in WM processes (Pavlov & Kotchoubey, 2022). Importantly, most prior studies utilized metrics such as averaged power, which give the impression that band activity is sustained and continuous over the length of long behavioral trials. As sustained average power can be the summation of transient, high-power oscillatory bursts (Jones, 2016), examining these transient burst events with trial-by-trial and non-averaged analyses may further advance WM models. In fact, Miller, Lundqvist and colleagues, using local field potential and spike recordings in non-human primates, have eloquently shown that oscillatory bursts (and not sustained activity) within the beta and gamma bands underlie WM processes within the PFC (Lundqvist et al., 2018; Lundqvist et al., 2016). Specifically, they found that the rate of gamma bursting increases during stimulus encoding (and subsequent readout or decoding) and decreases during WM delay, while beta bursting rates decrease during encoding and increase during WM delay. Further, beta bursting was negatively correlated with gamma bursting, with deviations in this beta/gamma rise and fall bursting pattern predicting behavioral errors.

While Miller and colleagues focused their empirical investigations within the beta band, the traditional alpha and beta-bands were grouped together (10-30 Hz) in their overarching model as both bands provide inhibition in different areas of cortex. This “push-pull” interplay between alpha/beta (10-30 Hz) and gamma (>30 Hz) bursting rate was conceptualized as alpha/beta bursts (primarily in cortical layers 5/6) carrying the control or inhibition of WM storage, and gamma bursts (primarily in cortical layers 2/3) carrying the encoding of sensory information (Buschman & Miller, 2022; Miller et al., 2018). Further, these authors have suggested that this alpha/beta versus gamma bursting reflects oscillatory states responsible for internal cognitive control and external sensory encoding, respectively (Lundqvist et al., 2024; Miller et al., 2018; Widge & Miller, 2019).

Building upon these burst findings in non-human primates and expanding the understanding of the potentially distinct roles of alpha and beta activity in human WM, in a recent MEG study Lundqvist and colleagues found that both alpha and beta burst rates were involved in human WM encoding, delay, and readout (Liljefors et al., 2023). Within occipital cortex, alpha and beta bursting decreased during stimulus presentation. Given the two bands showed differing temporal patterns and target versus distractor responses, the authors proposed beta burst rates support the transition from sensory processing to WM retention, while alpha burst rates suppressed unwanted sensory information. Further, they showed that prefrontal and parietal beta bursting increased before stimulus presentation, suggesting that beta suppressed retained information prior to target encoding (Liljefors et al., 2023). This is consistent with prior human work using averaged data that show alpha and beta power gate information flow, with alpha protecting against distractions, and beta flexibly activating task-relevant circuits (ElShafei et al., 2022; Zhou et al., 2023).

We have previously shown that bursting power, duration, and frequency span are critical burst features alongside burst rate that determine average power and correlate with human perception (Shin et al., 2017). However, most burst studies in WM to date have focused on burst rate, while the role of other burst features remains understudied (except (McKeon et al., 2023; Rodriguez-Larios & Haegens, 2023). One recent human EEG study expanded prior burst rate findings to amplitude, duration, and frequency span of beta bursts during WM across the scalp montage (Rodriguez-Larios & Haegens, 2023). Surprisingly, the prior Miller/Lundqvist findings of decreased beta burst rate during WM were not found, rather, beta burst amplitude and duration decreased during the WM delay, while burst peak frequency increased (Rodriguez-Larios & Haegens, 2023). Further, higher WM load was associated with decreased beta burst amplitude and duration, yet increased frequency and rate. These discrepancies and a still incomplete picture of burst features across different frequency bands (e.g., alpha, beta and gamma), networks (e.g., frontal and parietal), and WM stages (e.g., encoding vs maintenance) highlight the need for a more extensive examination of burst features to theorize the role of these rhythms in WM.

The objective of the current study was to identify the dynamic interplay between alpha/beta and high frequency activity (HFA; 55-80 Hz) bursts in human WM, with a more complete examination of various burst features (i.e., rate, amplitude, duration, frequency span) and error-related deviations in these features during both the encoding and maintenance of WM stimuli. We utilized a trial-by-trial burst characterization approach applied to a large, publicly available EEG dataset of 154 young adults collected during a Sternberg WM test. Results indicate that successful human WM is dependent on the rise and fall interplay between several alpha/beta and HFA burst features across the encoding and maintenance periods. Our extensive identification of bursting features involved in WM processes provides novel targets for treatment development in clinical populations with WM deficits.

## 2. METHODS

### 2.1 Participants

As previously described (Pavlov & Kotchoubey, 2020, 2021), 154 participants (82 female, mean age = 21.23, SD = 3.22) comprised the final sample. The participants had normal or corrected-to-normal vision and did not report any history of neurological or mental disease. All of them were Russian native speakers. The experimental protocol was approved by the Ural Federal University ethics committee.

### 2.2 Working Memory Task

As previously described (Pavlov & Kotchoubey, 2020, 2021), the task was a Sternberg working memory paradigm with Cyrillic alphabet letters with temporally distinct encoding and maintenance processing stages (see Figure 1A). The experiment entailed six different conditions: maintenance in memory of 5, 6 or 7 simultaneously presented letters in the alphabetical (manipulation task) or forward (retention task) order. In the retention task, the participants had to maintain in memory the original set as it was presented, and in the manipulation task, they had to, first, mentally reorganize the letters into the alphabetical order and then maintain the result in memory. After 6.7 s delay, a letter-digit probe appeared, and the participants indicated whether the probe was on the corresponding position either in the original set (retention task), or in the set resulted from the alphabetical reordering (manipulation task). Each of the six conditions (retention or manipulation of 5, 6 or 7 letters sets) entailed 20 consecutive trials. These six blocks of 20 trials were presented in a random order.

**Figure 1.**
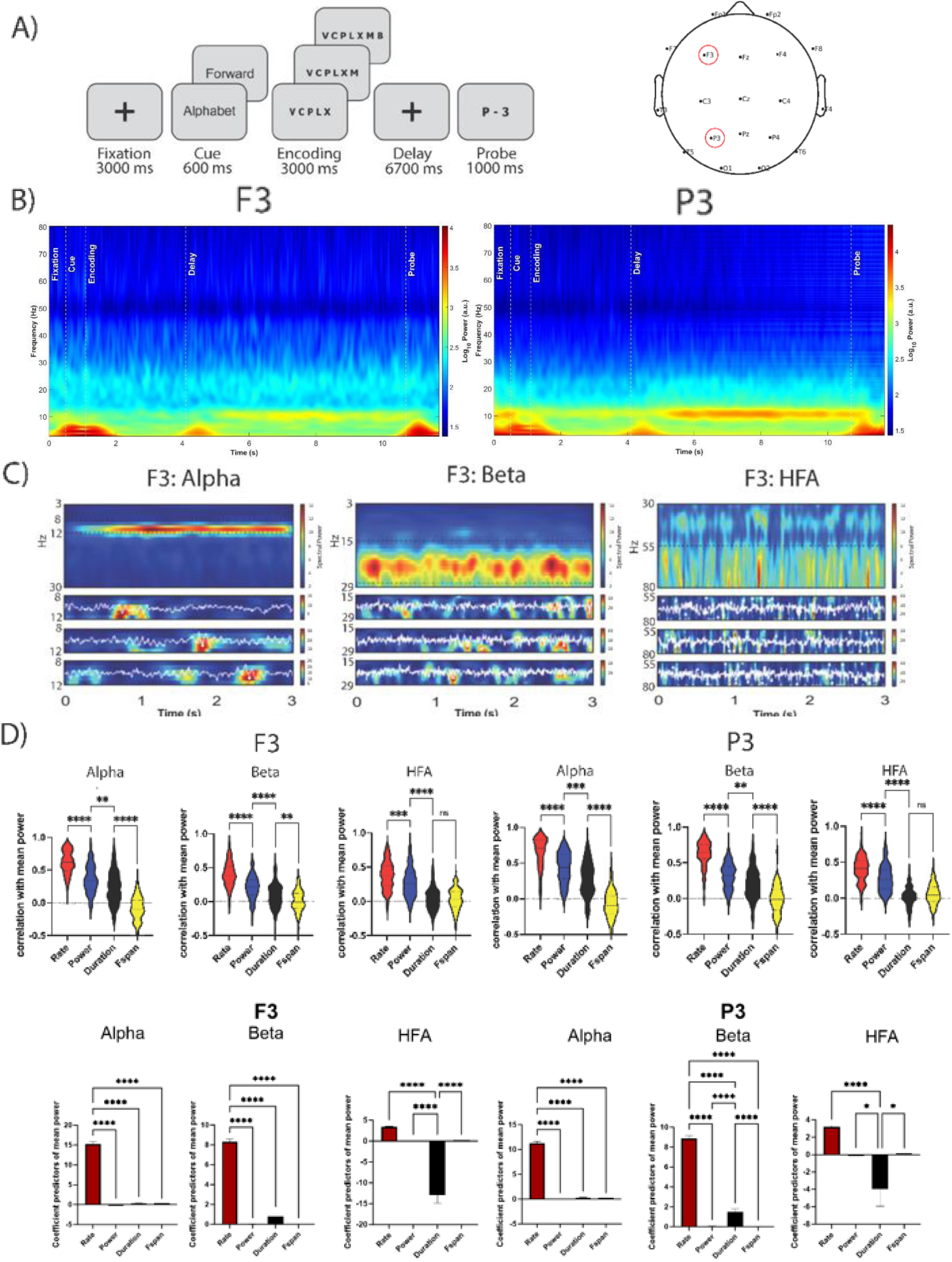
Initial spectrogram and sample burst raw data. A) Left. The experimental paradigm. Sets of Russian alphabet letters (5, 6 or 7) written in capitals were used as stimuli. An analogue using Latin letters and English words is shown (as noted by Pavlov et al., 2020). Duration of each phase of the task is indicated below. Right: EEG montage. B) Spectrogram across full trial for all participant/trials at F3 (left panel) and P3 (right panel). C) Sample averaged oscillatory power during individual recordings, decomposed into distinct alpha (left), beta (center), and HFA (right) bursts across three randomly sampled trials (electrode F3). D) Correlation analyses (top) and multiple regression analyses (bottom) between mean power and burst rate, burst power, burst duration, and burst frequency span.

### 2.3 EEG Recording & Preprocessing

As previously described (Pavlov & Kotchoubey, 2020, 2021), the EEG was recorded from 19 electrodes arranged according to the 10–20 system using Mitsar-EEG-202 amplifier with averaged earlobes reference. Two additional electrodes were used for horizontal and vertical EOG. EEG data were acquired with 500 Hz sampling rate and 150 Hz low-pass filter.

Pre-processing was conducted using the EEGLAB toolbox for MATLAB. Data was resampled to 250 Hz, 50 Hz line noise was removed with a notch filter (47-53 Hz), after which the data was high-pass filtered at 1 Hz, and low-pass filtered at 100 Hz. EEGLAB’s *clean_rawdata* function was used to remove channels for which a) signal was flat for 5+ seconds, b) high-frequency line noise was >3 standard deviations (SDs) above the mean, and c) there was < .85 correlation with nearby channels. This EEGLAAB function also removed data > 20 SDs outside the mean with the artifact subspace reconstruction algorithm and removed bad data periods with > 25% of channels out of acceptable range. Rejected channels (mean per participant = 2.3 [SD = 1.7]) were spherically interpolated and data was re-referenced to average. Independent component analysis (ICA) was conducted with the picard algorithm. Using EEGLAB’s *ICLabel*, components with a .8 or higher likelihood of being eye or muscle artifact were rejected.

To examine prefrontal and parietal regions, F3 and P3 electrodes (n = 1 interpolated; 99% non-interpolated) were extracted for further examination, as source localization was determined to be invalid given the 19-channel montage. Due to concerns about voltage conduction, multiple electrodes were not merged. F3 and P3 pairings were examined separately. Data were epoched to capture the full trial, including fixation cross (last 500 ms of 3000 ms window), cue (600 ms), stimulus encoding (3000 ms), and the working-memory maintenance or “delay” period (6700 ms). Baseline normalization was conducted using the fixation window. An epoch was rejected if it contained a value outside the +/- 150uV range. When examining discrete WM stages, we examined 500 ms of the fixation window (i.e., t = 2500-3000 ms of the full 3000-ms window), 2000 ms of the encoding window (i.e., t = 500-2500 ms of the 3000-ms window), and 4000 ms of the delay window (i.e., t = 1500-5500 ms of the 6700-ms window).

### 2.4 EEG Spectral Analysis

The time-frequency response (TFR) of each single-trial time series was calculated by convolution with a Morlet wavelet of the form

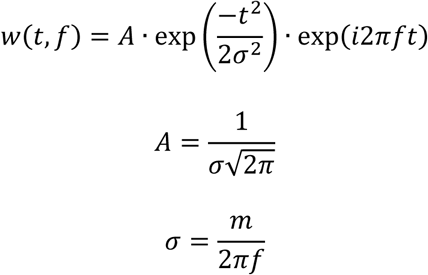

for each frequency of interest 𝑓 (3-80 Hz), and width 𝜎, which is determined by constant 𝑚 (selected here to be 7) that controls the number of cycles per wavelet. Visual inspection of the averaged TFR across all participants/trials throughout the task and across participants indicated presence of distinct alpha (8-12 Hz) and beta (15-29 Hz) band activity (Figure 1.B). Due to notable 50 Hz line noise, a notch filter was applied from 47-53 Hz. A clear pattern of bandlimited activity in the gamma range was not detected. However, based on prior work (Lundqvist et al., 2016) we remained motivated to investigate activity in the 55-80 Hz range of interest, but to clarify this was not clearly interneuron-mediated gamma oscillatory activity, we refer to activity in this range as “high frequency activity” (Iemi et al., 2022). Alpha, beta, and HFA bands were extracted for subsequent burst analysis.

### 2.5 Burst Analysis

Transient high-power “events” were detected and characterized using the SpectralEvents Toolbox (https://github.com/jonescompneurolab/SpectralEvents), which defines spectral events as any local maximum in the TFR above a power threshold within a user-defined band of interest. To be consistent with prior studies using similar methods, findMethod=1 was used as in Shin et al. (Shin et al., 2017) which is agnostic to event overlap, and the event threshold was set at 6x the median power (i.e., 6 factors-of-the-median [FOM]) across time and epochs for each frequency bin of the TFR (Levitt et al., 2020; Morris et al., 2023; Shin et al., 2017). Events were examined within the alpha (8-12 Hz), beta (15-29 Hz) and HFA (55-80 Hz) bands (Figure 1B-C). Each spectral event was characterized by its peak time/frequency within each trial, along with the event’s peak power, duration, and frequency span (f-span). Analysis was conducted on a subject-by-subject basis. *Event rate* was calculated by counting the number of events in the 2-s period of each epoch. *Event power* was calculated as the normalized FOM power value at each event maximum. The *Event duration* and *frequency span* (F-span) were calculated from the boundaries of the region containing power values greater than half the local maxima power, as the full-width-at-half-maximum in the time and frequency domain, respectively. Events with features greater than three standard deviations from the mean were removed. Z-scores of event features are reported (with baseline normalization using the fixation window).

To be consistent with prior studies that use a similar method of power thresholding to identify spectral bursts (i.e., “events”) in the time-frequency response of baseline neural activity (Kavanaugh et al., 2023; Levitt et al., 2020; Morris et al., 2023; Shin et al., 2017), we conducted all analyses here using a threshold of 6 factors-of-the-median (FOM). This cutoff of 6 FOM was originally chosen because it maximized the amount of trial-by-trial power variance explained by subthreshold versus suprathreshold activity (Shin et al., 2017). Thus, Shin et al. (2017) sought to obtain optimal sensitivity to high-power time-frequency fluctuations while minimizing the bias induced by the presence (or lack thereof) of a constant rhythm in the band-of-interest (BOI). To verify that 6 FOM was a reasonable choice for the present study and each frequency band, we quantified the across-trial Pearson’s correlation between mean power within the BOI and percent area above the cutoff at various threshold values ranging from 0.25-16 FOM (**Supplemental Figure 1**). We found that the correlation coefficient averaged across participants peaked near 6.0 FOM in each frequency band and electrode, demonstrating that this value was optimal for identifying spectral bursts. Further, we estimated probability of observing a spectral burst as a function of threshold (**Supplemental Figure 1**) and its corresponding cumulative distribution function (CDF). 1-CDF in Figure 2 shows the proportion of total possible bursts above the cutoff.

**Figure 2.**
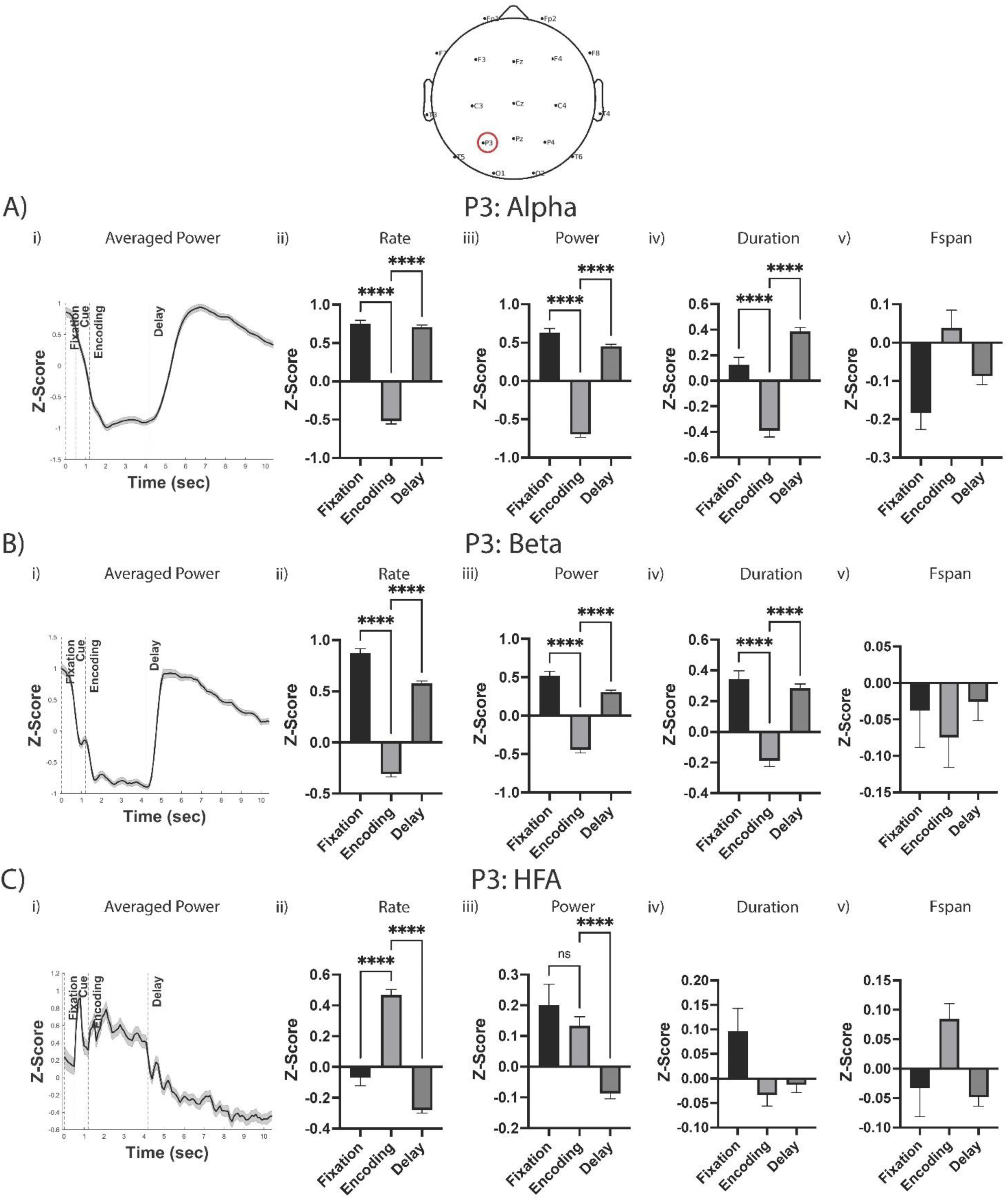
Burst power, duration, and frequency span, not just burst rate, change across WM stages in a pattern reflective of averaged power at P3. i) Averaged power time series across trial. ii-v) Oscillatory bursting patterns across fixation, encoding, and delay stages for burst rate (ii), burst power (iii), burst duration (iv), and burst frequency span (v) within alpha (A), beta (B), and HFA (C) bands. Asterisks denote statistical significance. Pairwise comparisons only displayed when overall model was statistically significant after Bonferroni correction.

### 2.6 Statistical Analyses

Statistical analyses were conducted in Matlab-2022a and GraphPad Prism 10. A series of Pearson correlation analyses examined the association between TFR and event feature correlates within each band at each time point, with one-way analyses of variance (ANOVA) examining the differences between event features on averaged power across participants (with Tukey’s correction for pairwise comparisons). To simultaneously examine the contribution of each burst feature to average power, a series of regression analyses were conducted with burst features as independent variables and average power as the dependent variable. After showing appropriate correlation between burst features (i.e., r ≤ .5; Figure 7), a regression analysis was conducted at each time point and then an ANOVA compared the averaged regression coefficient (across all timepoints) of each feature (ANOVA Bonferroni corrected for two electrodes and three bands, i.e., raw p-value multiplied by 6; Tukey correction for pairwise comparisons).

Two-way repeated measures analysis of variance (2-RM-ANOVA) was utilized for most analyses, although if missing data was present, a mixed-effects model (i.e., restricted maximum likelihood [REML]) was alternatively conducted based on the default parameters in GraphPad to fit a mixed effect model to repeated measures data. Data were entered in a grouped format, within which each participant (set as random factor) had data for each relevant time point (set as fixed factor) and group (i.e., correct versus incorrect or F3 versus P3; set as fixed factor). Missing data was predominantly present only when bursting was not detected for that participant/timepoint, so the burst rate = 0 but burst features were unavailable (as there was no burst to measure features). The primary effect of time (i.e., fixation, encoding, delay) and the time x accuracy interaction were examined for relevant analyses (for analyses comparing regions, i.e., Figure 3, the time x region interaction was examined). Pairwise comparisons after models were corrected with Sidak correction. The primary effect of time and the time x accuracy interactions are displayed separately in figures to improve readability.

**Figure 3.**
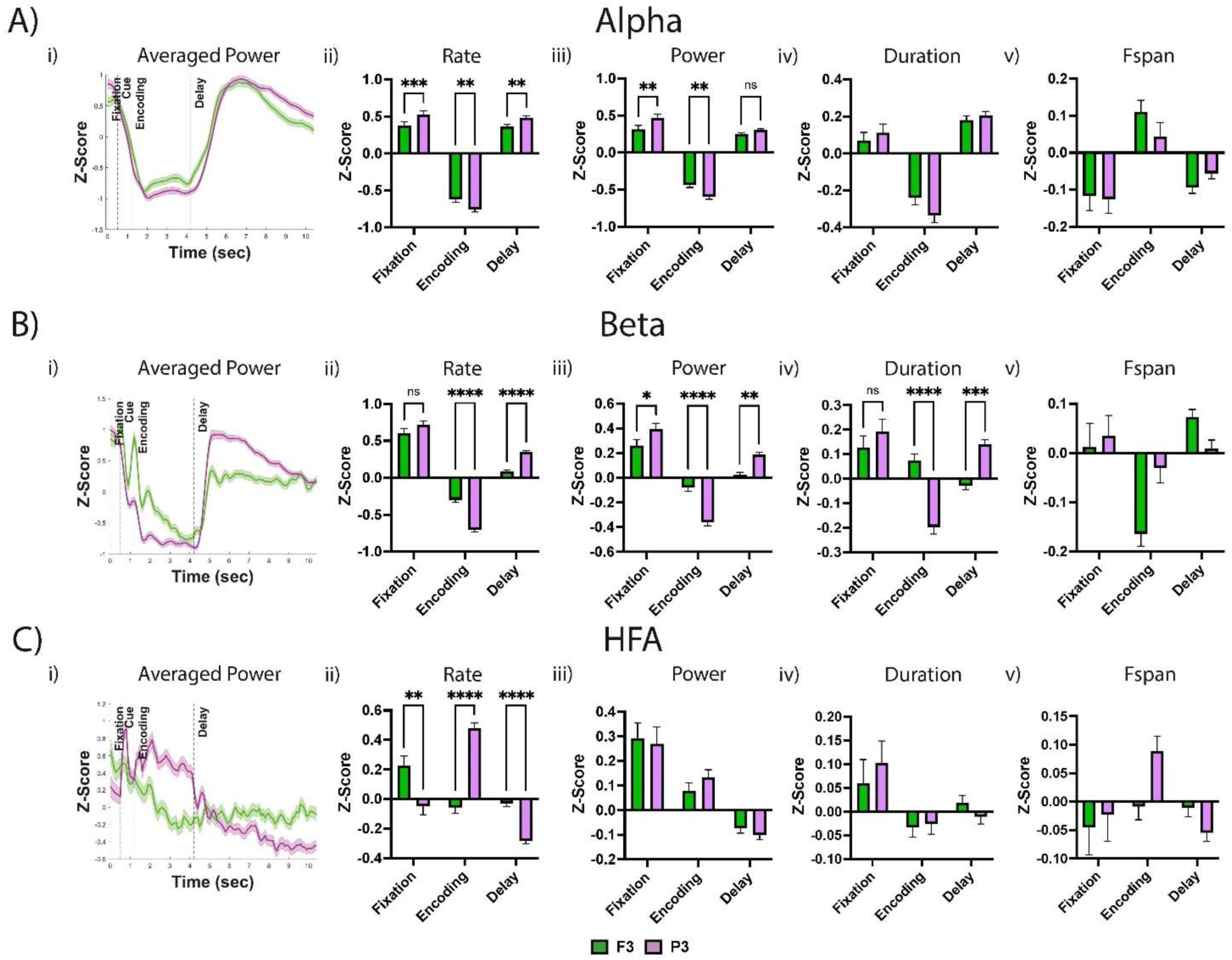
Frontal versus parietal recoding was associated with diminished bursting patterns across alpha, beta, and HFA bands. i) Averaged power time series across trial for F3 and P3 electrodes. ii-v) Time x electrode analyses examining differences between electrodes in oscillatory bursting patterns across fixation, encoding, and delay stages for burst rate (ii), burst power (iii), burst duration (iv), and burst frequency span (v) within alpha (A), beta (B), and HFA (C) bands. Asterisks denote statistical significance. Pairwise comparisons only displayed when overall model was statistically significant after Bonferroni correction.

Bonferroni correction was applied to all RM-ANOVA and mixed model analyses to account for two electrodes (F3 & P3), bands (alpha, beta, HFA) and event features (rate, power, duration, frequency span; i.e., raw p-value multiplied by 24). Multiple comparison corrected p-values are reported (e.g., uncorrected p-value multiplied by 24, unless otherwise noted), with statistical significance set at p < .05. If a given set of analyses were not Bonferroni corrected, the “uncorrected p” term will be used (e.g., follow-up analyses). Raw scores converted to z-scores (with baseline normalization using the fixation window) were utilized for analyses unless indicated otherwise. Only correct trials were included, unless mentioned otherwise.

To simultaneously examine the contribution of each burst feature to total task accuracy, a series of regression analyses were conducted with burst features as independent variables and task accuracy as the dependent variable. Analyses were conducted separately for the Encoding and Delay phases. After showing appropriate correlation between burst features (i.e., r ≤ .5; Figure 7), a regression analysis was conducted at each time point and then an ANOVA compared the averaged regression coefficient (across phase timepoints) of each feature (ANOVA Bonferroni corrected for two electrodes and three bands, i.e., raw p-value multiplied by 6; Tukey correction for pairwise comparisons).

Features that were more strongly associated with task accuracy were further examined across the EEG scalp montage. A similar series of linear regression analyses (burst features as independent variables) was conducted at each electrode (for each time point). The burst rate coefficient was extracted for alpha and beta band analyses and the burst duration coefficient was extracted for HFA band analyses, and these coefficients were examined across electrode regions (Anterior: FP1/2, F3/4/z/7/8; Central: C3/z/5, T3/4; Posterior: T5/6, P3/z/5, O1/2). Single-sample t-tests compared the coefficient of each region to zero (Bonferroni corrected for three regions and two time points [i.e., raw p-value multiplied by 6]). Similarly, and to build upon the HFA – alpha/beta contrast findings, we then conducted the same RM-ANOVA of this contrast score across all electrodes, extracted the model F statistic of each electrode’s analysis, and examined across brain regions with a one-sample t-test (Bonferroni corrected for three regions and two time points [i.e., raw p-value multiplied by 6]).

Finally, we examined the possible differential effect of bursting dynamics on task accuracy for manipulation versus retention WM demands and low (5 items) versus medium (6 items) versus high (7 items) WM load by conducting similar regression models, extracting regression coefficients, and comparing between WM demands with paired sample t-tests (no correction applied; p < .05).

## 3. RESULTS

Participants completed a version of the Sternberg working-memory paradigm (**Figure 1.A**), within which they were presented and asked to manipulate or retain a set of visually-presented alphabet letters (Pavlov & Kotchoubey, 2020, 2021), while EEG was continuously recorded. We focused only on activity from EEG sensors above frontal (electrode F3) and parietal (P3) cortex, to relate results to prior studies. Examination of non-averaged individual trials shows bursts of high power in each band (**Figure 1.C**). The rate, power, duration, and frequency span of oscillatory bursts were examined within these bands in n = 154 participants. Low frequency, transient activity during stimuli presentation/removal appeared to reflect evoked responses, and therefore we continued to focus our predetermined analyses in alpha, beta, and HFA bands.

### 3.1 Burst rate is most strongly correlated with trial averaged power

To investigate what features of the burst patterns underlie averaged power metrics, we examined the correlation between the average of each burst feature (i.e., rate, power, duration, frequency span) and the averaged power across the trials. Across all bands and electrodes, burst rate was always the most strongly associated with averaged power, followed by burst power, burst duration, and burst frequency span, with statistically significant differences between these features detected in pairwise follow-up analyses for the majority of bands/features (**Figure 1.D.Top panel**; Alpha F3: F = 309, p < .0001; Alpha P3: F = 316, p < .0001; Beta F3: F = 158, p < .0001; Beta P3: F = 299, p < .0001; HFA F3: F = 181, p < .0001; HFA P3: F = 196, p < .0001). In examining notable correlations between features, frequency span had a consistently positive association with rate and a negative association with duration (**Figure 7**). We then examined if specific burst features were most strongly associated with averaged power than others via multiple regression with all features loaded in each model. The burst rate was the feature most strongly associated with average power across all bands and electrodes (**Figure 1.D.Bottom panel**; F3 – Alpha: F = 592 p < .0001; F3 – Beta: F = 340, p < .0001, F3 – HFA: F = 54.7, p < .0001; P3 – Alpha: F = 635, p < .0001; P3 – Beta: F = 468, p < .0001; P3 – HFA: F = 8.96, p < .0001).

### 3.2 Average power change across WM stages reflects changes in burst features

Consistent with prior studies on averaged alpha/beta power during WM (Paulo et al., 2023; Pavlov & Kotchoubey, 2020, 2021; Roux et al., 2012), averaged alpha and beta power at P3 decreased during the fixation cue and increased during the delay, while averaged HFA power increased during the fixation cue and decreased during the delay (**Figure 2.A-C.i**). Given that on individual trials alpha/beta and HFA power emerge as transient bursts, we next examined which burst features (rate, power, duration, and frequency span) contributed to the observed average power differences.

Alpha and beta exhibited a similar pattern of bursting across WM stages (**Figure 2.A-B.ii-v**). Specifically, the alpha/beta burst rate, burst power, and burst duration reflected the averaged power pattern of a decrease in activity from fixation to encoding and an increase from encoding to the delay (Alpha burst rate: F = 408; p < .0001; Alpha burst power: F = 245; p < .0001, Alpha burst duration: F = 72.5, p < .0001; Beta burst rate: F = 418; p < .0001; Beta burst power: F = 138, p < .0001; Beta burst duration F = 48.1, p < .0001). There was no significant pattern in the frequency span of alpha or beta bursts (both p > .05).

The rate and power of HFA bursting largely reflected the averaged power pattern, in that the rate and power of HFA bursting displayed the inverse pattern of alpha/beta bursting (**Figure 2.C.ii-v;** HFA burst rate: F = 111; p < .0001; HFA burst power: F = 15.2, p < .0001). The rate of HFA bursting increased during encoding and decreased during the delay, while the power of HFA bursts was characterized by a decrease during the delay. There was no significant pattern in the duration or frequency of HFA bursting (both p > .05).

<u>Taken together</u>, alpha/beta and HFA bursting follow inverse patterns of rising and falling across WM stages at P3 (but not at F3). Consistent with prior work, this pattern was found in burst rate (Liljefors et al., 2023; Lundqvist et al., 2018; Lundqvist et al., 2016) and in alpha/beta burst features (Rodriguez-Larios & Haegens, 2023), but uniquely, we found this pattern was also present in the power and duration of HFA bursts.

### 3.3 Dynamic variation in bursting differs between parietal and prefrontal electrodes

Given the distinct roles of prefrontal versus parietal cortex in WM processes (Crowe et al., 2013; Murray et al., 2017), we then examined activity at the F3 electrode and compared to P3 electrode findings (**Figure 3**).

The overall trends in the alpha and beta dynamics across the WM stages were consistent across electrodes, with a few notable differences suggesting overall enhanced dynamic variation in parietal (P3) compared to prefrontal (F3) sites (**Figure 3.A-B**). Similar to P3, at F3 the averaged alpha and beta power decreased from the fixation to encoding period and increased from encoding to delay, and these effects were mainly driven by corresponding changes in burst rate, power and duration (results were mixed for frequency span, see **Supplemental Table 1** for significance across WM stages).

Notable differences across electrodes include that the averaged alpha/beta power decrease from fixation to encoding was faster and stronger in the P3 electrode, as was the increase from the encoding to delay period, and that these effects were greater in the beta band (**Figure 3**). These differences were driven by changes in burst rate and power in both bands, and additionally duration differences in the beta band (**Figure 3.A-B**; Alpha Rate: F = 16.0, p < .0001; Alpha Power: 13.1, p < .0001; Beta Rate: F = 27.4, p < .0001; Beta Power: F = 29.4, p < .0001; Beta Duration: F = 25.8, p < .0001).

The averaged HFA power dynamics across WM stages was markedly different at F3 and P3 and emerged solely from significant differences in HFA rate dynamics. Frontal activity at F3 was characterized by a decrease during encoding and plateau during the delay, while at P3, HFA increased during encoding and decreased during the delay (**Figure 3.C.i**). HFA burst rates were higher at P3 during encoding and higher at F3 during the delay (F = 45.2, p < .0001; **Figure 3.C.ii**).

<u>Taken together</u> with prior theories suggesting different roles of parietal and frontal cortex in sensory encoding and cognitive control (Crowe et al., 2013; Murray et al., 2017), our results suggest that the fast alpha/beta changes at P3 may reflect earlier sensory encoding in this region while the distinct differences in HFA band activity across electrodes may more directly reflect differences in the role of these parietal and frontal cortex in sensory versus cognitive control.

In examining the correlation between band/features across all timepoints, there was a negative association between HFA burst rate and alpha/beta burst features at P3 **(Supplemental Figure 4.A**; beta burst rate: r = -.16; p < .0001; beta burst power: r = -.10; p < .0001; alpha burst rate: r = -.22; p < .0001; alpha burst power: r = -.18; p < .0001; alpha burst duration: r = -.11; p < .0001). Further, we found that alpha/beta and HFA had mirrored patterns across timepoints at P3 (**Supplemental Figure 4.B**). Interestingly, this negative association was not found at F3; rather, there was a positive association between beta rate and HFA rate at F3 (**Supplemental Figure 4.A**; r = .11; p < .0001).

<u>Taken together</u>, we confirmed non-human primate findings that alpha/beta and gamma/HFA bursting show inverse patterns (although at P3, not F3) for differing WM demands (Lundqvist et al., 2018; Lundqvist et al., 2016), and uniquely found that this association also include the power and duration of bursts.

### 3.4 Incorrect responses are associated with weaker variation in burst features across WM stages

We then examined differences in burst patterns between correct and incorrect trials (mean number of incorrect trials= 21.2 [SD = 7.9]; range 4-42). At P3, incorrect trials had weaker dynamic variation (i.e., less pronounced variation in bursting) across WM stages compared to correct trials (**Figure 4**). Correct-incorrect differences were observed in the burst rate for alpha, beta, and HFA bands and additionally in power in the beta band (**Figure 4.A-C.ii**; Alpha Rate: F = 14.2, p < .0001; Beta Rate: F = 21.5; p < .0001; Beta Power: F = 11.2, p < .0001; HFA Rate: F = 6.3; p = .048). Findings were overall less robust at F3 (**Supplemental Figure 3**) but followed a qualitatively similar pattern to P3. A comparison of the difference between alpha/beta and HFA burst rate dynamics across WM stages (i.e., HFA rate – combined alpha/beta rate) also showed that weaker variation was indicative of incorrect responses (**Figure 5.A.i**), suggesting that a strong inverse relationship between HFA and alpha/beta is an important component of accurate working-memory processing (Miller et al., 2018; **Figure 5.A.i**; Rate: F = 24.4, p = 3.1e-05; Power: F = 8.8, p > .05; Duration: F = 3.8, p > .05; Frequency Span: F = .23, p > .05).

**Figure 4.**
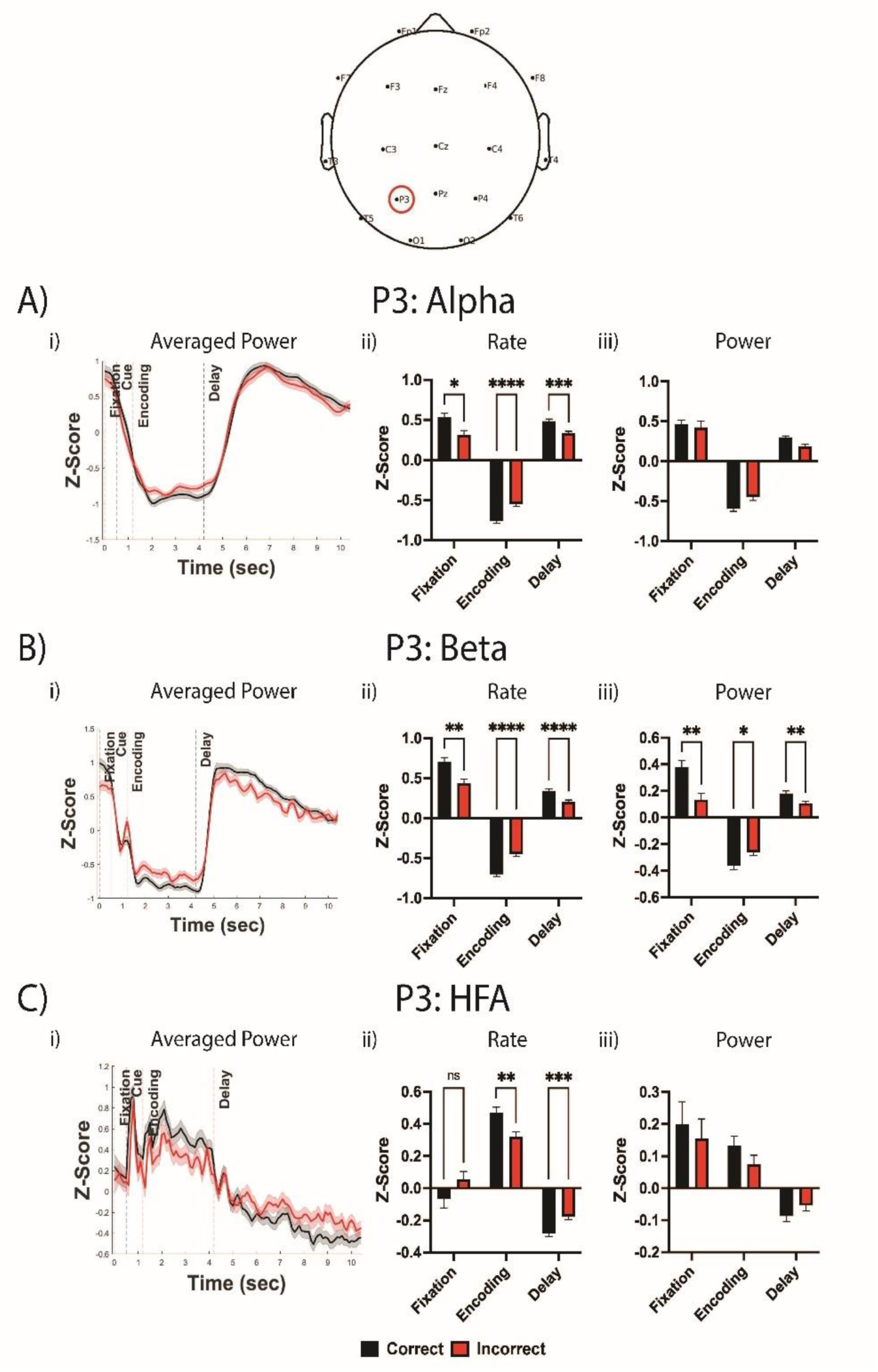
Incorrect responses were associated with diminished parietal bursting patterns across alpha, beta, and HFA bands. i) Averaged power time series across trial for correct and incorrect trials. ii-v) Time x accuracy analyses examining differences between electrodes in oscillatory bursting patterns across fixation, encoding, and delay stages for burst rate (ii) and burst power (iii) within alpha (A), beta (B), and HFA (C) bands. Asterisks denote statistical significance. Pairwise comparisons only performed when overall model was statistically significant after Bonferroni correction.

**Figure 5.**
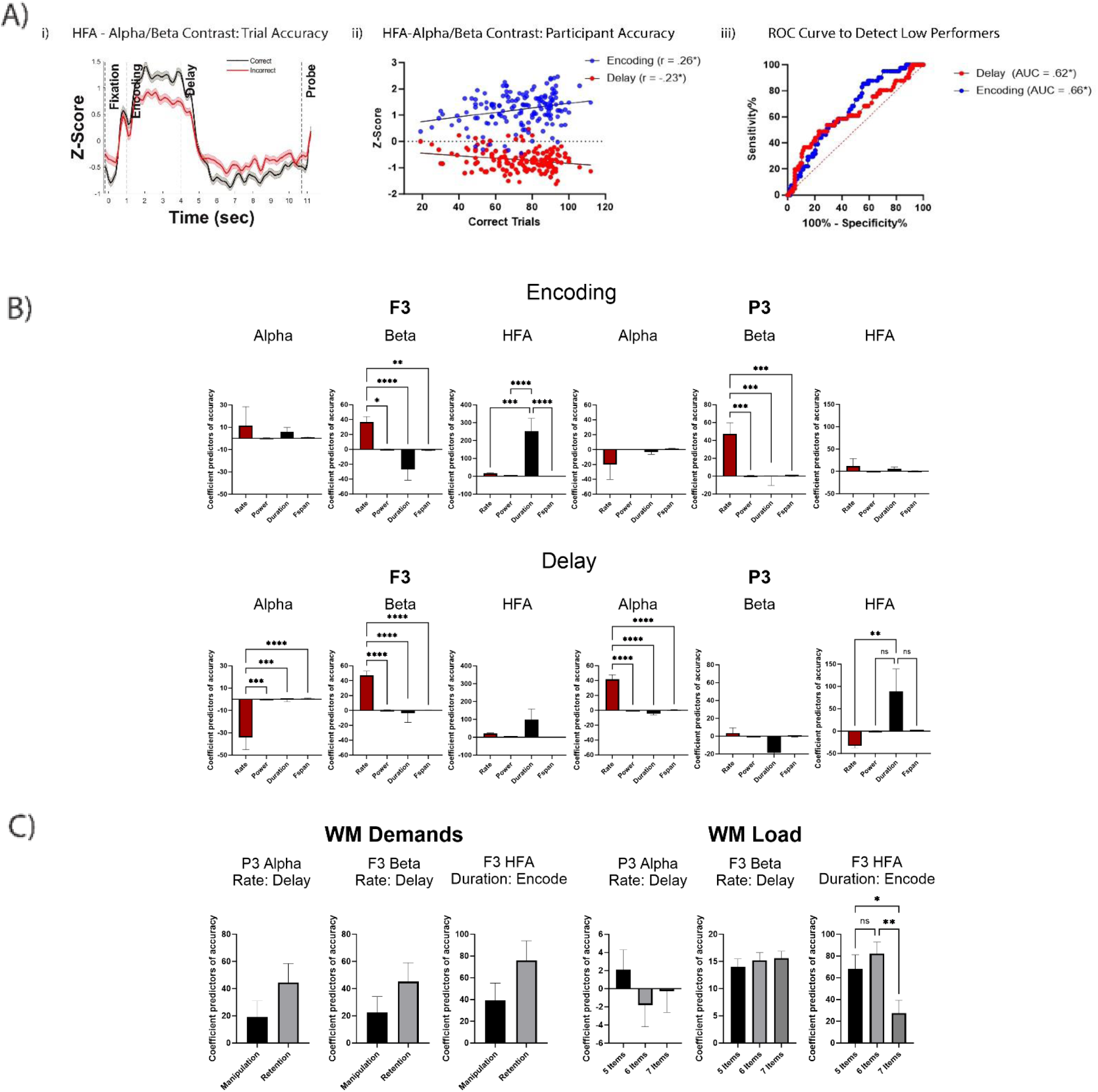
A. i) Time x accuracy analysis examining the difference between correct and incorrect trials in the contrast between HFA and alpha/beta bursting rate. ii) Correlation between the HFA-alpha/beta bursting contrast during encoding and delay and total participant task accuracy. iii) Receiver operating characteristic curve of the HFA-alpha/beta contrast in differentiating low performing participants from the rest of the sample. B. Multiple regression models and follow-up ANOVAs examining burst feature predictors of total WM task accuracy, examined across electrodes, bands, and the encoding (top) and delay (bottom) periods. C. Multiple regression models and ANOVAs examining burst feature predictors of total WM task accuracy across differing WM demands/load.

In examining overall participant performance, there was a differential association between this HFA – alpha/beta contrast (i.e., “HFA-ABC”) score and overall participant task accuracy, with a positive association found during stimulus encoding and a negative association found during the WM delay (**Figure 5.A.ii**; encoding: r = .26, uncorrected p < .05; delay: r = - .23, uncorrected p < .05). Further, in dividing the sample into the lower 25th percentile (i.e., < 25^th^ percentile) and rest of the sample (i.e., ≥ 25^th^ percentile) to detect low performers, a receiver operating curve (ROC) of this HFA – alpha/beta contrast score was able to adequately differentiate the low performers from the rest of the sample (**Figure 5.A.iii**; delay: area under the curve [AUC] = .62, uncorrected p < .05; encoding: AUC = .66, uncorrected p < .05).

<u>Taken together</u>, building upon prior work in frontal cortex of non-human primates (Lundqvist et al., 2018; Lundqvist et al., 2016), we found that incorrect trials were associated with reduced dynamic variation of burst patterns in frontal and parietal cortex, and that this pattern was most notable in parietal cortex. Furthermore, reduced alpha/beta-HFA dynamics in parietal cortex were associated with poor overall WM task performance.

### 3.5 Alpha/beta bursting rate and HFA bursting duration relate to task accuracy

We then examined if specific burst features were most strongly associated with task accuracy than others (**Figure 5.B**). Within the alpha band, the burst rate at P3 during the delay was most strongly associated with total accuracy, while the burst rate at F3 during the delay was most negatively correlated with total accuracy (F3 – Delay: F = 9.9, p < .0001; P3 – Delay: F = 49, p < .0001). Within the beta band, the burst rate at F3 and P3 during encoding and at F3 during the delay was most strongly associated with total accuracy (F3 – Encode: F = 10, p < .0001; F3 – Delay: F = 12, p < .0001; P3 – Encode: F = 8.8, p < .0001). Within the HFA band, the burst duration at F3 during encoding and P3 during the delay was most strongly associated with total accuracy (F3 – Encode: F = 11, p < .0001); P3 – Delay: F = 4.5, p = .029).

Next, we examined the metrics with strong associations to task accuracy across the EEG montage (**Figure 6**). The rate of beta bursting was most strongly associated with overall task accuracy in posterior brain regions during encoding (t = 4.6, p = .02), while the beta burst rate was most strongly associated with overall task accuracy in anterior brain regions during the delay (t = 3.99, p = .04). No patterns were observed in alpha burst rate or HFA burst duration. Further, to also examine HFA-ABC across the montage in a similar manner, the difference between correct and incorrect trials for the HFA-ABC was most substantial in parietal brain regions during both encoding (t = 7.6, p = .002) and the delay (t = 6.6, p = .004). Full electrode montage analyses were largely consistent with initial single electrode findings.

**Figure 6.**
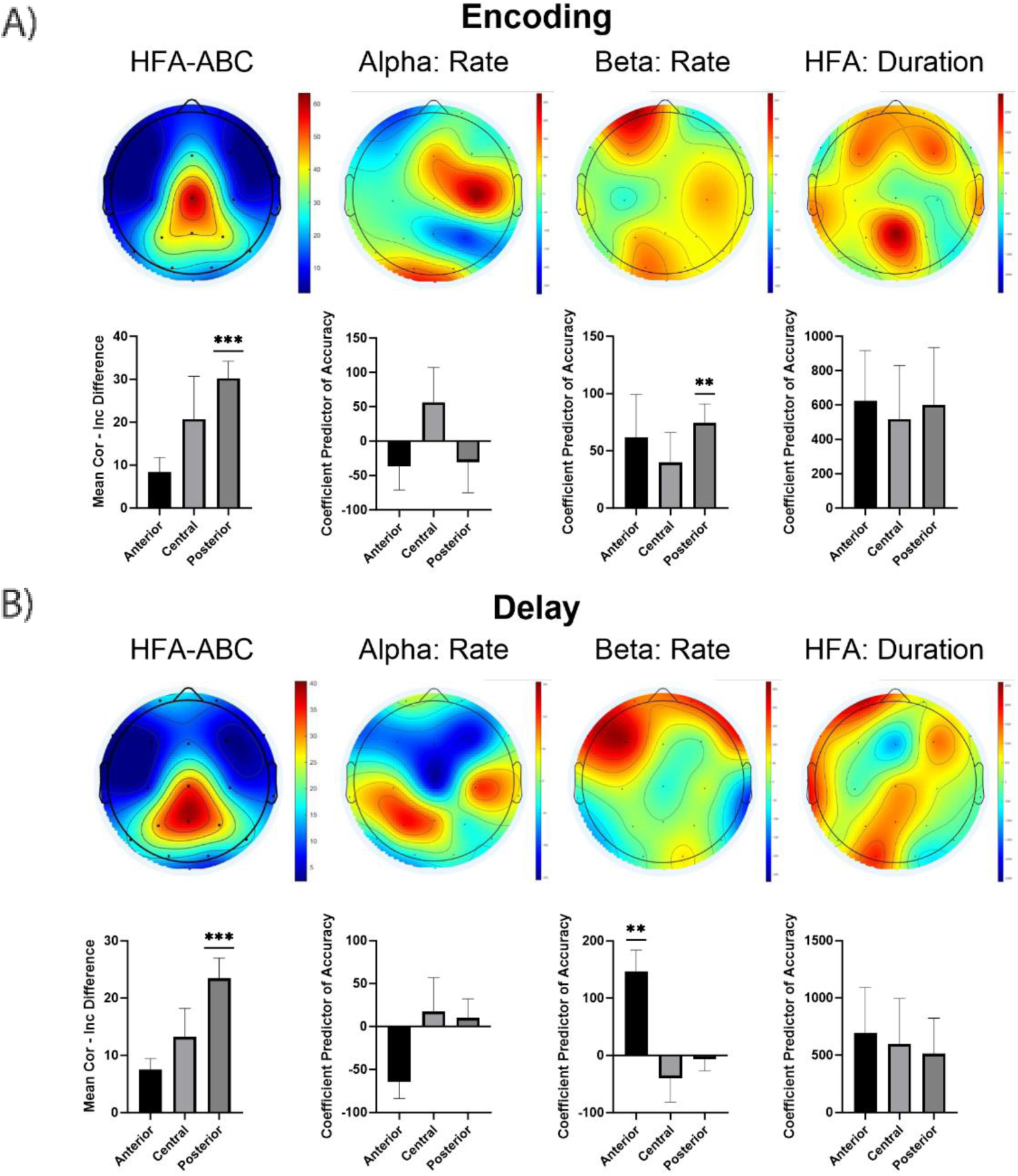
Spatial distribution of burst features during encoding (A) and delay (B) periods, as shown by EEG montage (top) and bar graphs separating different montage regions (bottom; Anterior: FP1/2, F3/4/z/7/8; Central: C3/z/5, T3/4; Posterior: T5/6, P3/z/5, O1/2).

**Figure 7.**
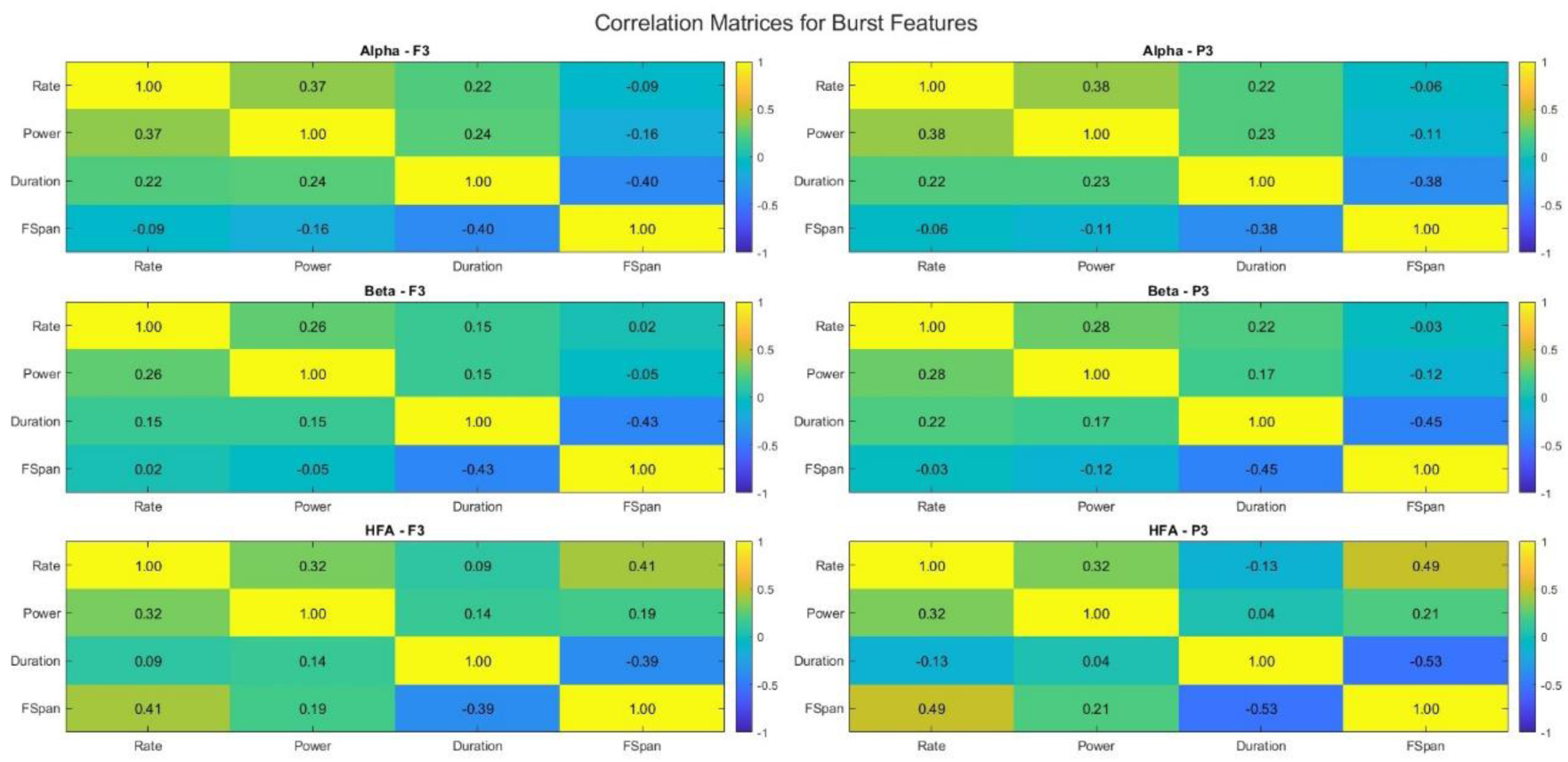
Correlation matrices of burst features across bands and electrodes.

We finally examined these findings across WM demands (i.e., manipulation versus retention) and WM load (i.e., low versus medium versus high). The relationship between bursting and task accuracy was not significantly different during manipulation versus retention demands (**Figure 5.C**; all uncorrected p values > .05). No findings for WM load were observed in alpha and beta bands, although there were HFA findings. Specifically, the weakest relationship between task accuracy and HFA bursting at F3 during encoding was found during high WM load compared to low and medium WM load (F = 5.8, uncorrected p = .004). To provide behavioral performance context to these demand findings, as a group, participants performed significantly better during retention demands compared to manipulation demands (paired sample t-test: t = 16.5, p < .0001), and performed steadily worse with more items (ANOVA; F (2,459) = 77.5, p < .0001; pairwise comparisons: 5 items > 6 items > 7 items).

## 4. DISCUSSION

Utilizing a trial-by-trial burst characterization approach within a large, publicly available EEG-WM dataset, we demonstrated the interplay between alpha/beta and high frequency activity (HFA; 55-80 Hz) bursts in human WM. Averaged power dynamics were driven by oscillatory bursts across bands, most consistently reflected in burst rate. Alpha/beta and HFA bursts displayed complementary roles in WM processes, in that alpha and beta bursting decreased during encoding and increased during delay, while HFA bursting had the opposite pattern. Weaker changes in this pattern across WM states were associated with incorrect responses, such that incorrect trials had less dynamic variation in alpha/beta and HFA burst dynamics. This reduced dynamic variation was also associated with poor overall task performance, and together, suggests reduced alpha/beta-HFA dynamics may reflect both a trial-by-trial and overall performance level marker of WM deficits (Results are summarized in Supplementary Table 1.)

Although oscillatory burst rate is the most commonly utilized burst metric in the literature, prior work has identified the critical importance of measuring burst power, duration, and frequency span to most precisely understand underlying neocortical circuit origins of averaged power signals during response inhibition (Enz et al., 2021), perception (Sherman et al., 2016; Shin et al., 2017), and working memory (McKeon et al., 2023; Rodriguez-Larios & Haegens, 2023). While several burst features were associated with averaged power and displayed temporally similar patterns to averaged power, the burst rate across alpha, beta, and HFA bands was most strongly associated with averaged power during WM in multiple regression models. The duration of HFA bursting was most negatively associated with averaged power, likely reflecting that more bursts (that are shorter in duration) leads to a larger averaged power, compared to bursts that are longer duration (but are fewer in amount). This is consistent with our prior work on beta burst rate in perception (Shin et al., 2017), and gamma burst rate and power in WM (McKeon et al., 2023). Similarly, the burst rate, power, and duration generally reflected the averaged power time course pattern across WM demands. Although frequency span was the least correlated to averaged power and did not consistently reflect the time course pattern in this study, it has shown utility as an important clinical marker in other contexts (Morris et al., 2023). Further, frequency span was positively correlated with rate and negatively correlated with duration, consistent with our prior findings (Shin et al., 2017). Broadly, our results highlight the continued utility of moving away from averaged power metrics and favor a measurement approach which captures the dynamics of oscillatory bursts in WM.

Our findings that the rate, power, and duration of alpha and beta bursts consistently decreased during stimulus encoding and increased during WM delay, is highly consistent with prior oscillatory burst rate findings (Liljefors et al., 2023; Lundqvist et al., 2018; Lundqvist et al., 2016) and averaged power findings (Paulo et al., 2023; Pavlov & Kotchoubey, 2020, 2021; Roux et al., 2012). Time course pattern differences were present between alpha and beta bursting, as beta sharply increased bursting during the delay, while alpha increased more slowly, consistent with recent work on the nuanced differences between alpha and beta bursting in WM (Liljefors et al., 2023). Our findings are complementary to recent findings that compared the WM delay to fixation using a different approach to burst characterization, and found that the WM delay was associated with lower beta burst amplitude, lower burst duration, and higher burst frequency, although with no change in burst rate (Rodriguez-Larios & Haegens, 2023).

While alpha/beta findings were largely consistent across F3 and P3 electrodes, changes across WM stages occurred earlier and were stronger at P3 than at F3, particularly in the beta band, consistent with the interpretation that parietal cortex represents fast sensory encoding and frontal cortex more higher-level processing (Crowe et al., 2013; Murray et al., 2017). In contrast, parietal HFA bursting rate increased during encoding and decreased during the delay, while frontal HFA burst rate peaked at fixation, decreased during encoding, and was stable during the delay. Importantly, note that we describe the 55-80 Hz range in our study as high frequency activity (HFA), instead of gamma, as a pattern of bandlimited activity was not detected, and it was unclear whether observed activity reflected interneuron-mediated gamma oscillations. We compare current HFA findings and prior gamma-related findings despite their potential differences, as both are similarly associated with neuronal excitability, while alpha/beta activity are associated with neuronal inhibition (Iemi et al., 2022; Miller et al., 2018; Wang, 2010).

Parietal gamma bursting rate was consistent with prior burst patterns within the PFC of non-human primates (Lundqvist et al., 2018; Lundqvist et al., 2016). Our frontal versus parietal HFA finding is highly consistent with prior work, which similarly found that gamma (60-80 Hz) power strongly increased during encoding and decreased during the WM delay in (right) parietal cortex, while gamma power in left frontal cortex increased to a lesser degree and remained relatively stable during the WM delay (Roux et al., 2012). Further, increased gamma-related activity at stimulus encoding has been shown to be more pronounced in parietal cortex compared to prefrontal cortex (Murray et al., 2017). It is possible these frontal versus parietal differences reflect different WM stages of processing, in that frontal gamma band activity peaks during the fixation as task demands are assessed and/or filtered, but parietal gamma band activity peaks at stimulus presentation to receive the sensory input, consistent with the known role of frontal versus parietal networks (Crowe et al., 2013; Murray et al., 2017).

Most notably, this is the first human study to our knowledge to show an anti-correlated and mirrored pattern between the beta and gamma/HFA burst rate during WM (although only at P3 electrode and not the F3 electrode), and replicates the highly influential work of Miller and Lundqvist that showed this pattern within single PFC neurons of non-human primates (Lundqvist et al., 2018; Lundqvist et al., 2016). It is unclear why prior frontal findings were not replicated, but methodological limitations related to the signal-to-noise ratio in scalp EEG cannot be ruled out. Further, it is the first study in either humans or non-human primates to show that this pattern extends into an alpha-HFA pattern, and extends beyond burst rate to also include burst power and duration. Consistent with its known cognitive control capabilities outside of WM (Enz et al., 2021; Wessel, 2020), alpha/beta has been conceptualized as a cognitive control mechanism or state during the WM process. During WM, alpha/beta activity has been found to gate information flow and recruit task-relevant circuits (ElShafei et al., 2022; Zhou et al., 2023), while gamma has been found to process perceptual information and maintain that information during the delay (Lundqvist et al., 2016; McKeon et al., 2023). This alpha/beta cognitive control state is conceptualized as being activated during task initiation (i.e., beta activity rises during fixation), then deactivated during stimulus presentation to allow for perceptual encoding (i.e., beta drops and gamma rises during encoding), and then re-activated during the delay (i.e., beta rises and gamma drops) as cognitive control demands are critical when ‘remembering’ the stimuli (Miller et al., 2018).

We showed for the first time in humans that weaker variations in these patterns across WM stages were associated with incorrect responding. Such deviations were primarily restricted to burst rate, although parietal beta burst power was also implicated (and not frontal HFA burst rate). A unified pattern emerged across bands and electrodes, in that the rise and fall of the burst rate across WM stages in correct trials was not as pronounced in incorrect trials. Incorrect trials had a lower burst rate when a higher burst rate was expected (i.e., not enough bursting) and a higher rate when a lower rate was expected (i.e., too much bursting). One interpretation is that, during encoding, incorrect trials had too many alpha/beta bursts and not enough HFA bursts, while during the delay, incorrect trials had too many HFA bursts and not enough alpha/beta bursts. A more nuanced finding was previously shown in monkeys, in that during incorrect trials, the beta/gamma bursting pattern reflected the expected pattern for the incorrect response (i.e., not the pattern expected for the correct response; (Lundqvist et al., 2018). Further, we found that correct trials had a more pronounced difference between HFA and alpha/beta bursting (i.e., HFA-alpha/beta contrast [“HFA-ABC”]) than incorrect trials, in that correct trials were associated with higher HFA and lower alpha/beta during encoding and the opposite pattern during the delay.

We next examined correlates of overall task performance. This HFA – ABC strongly correlated to overall participant accuracy and differentiated low performing participants from the rest of the sample. When comparing which features were most strongly associated with task accuracy, we found that the beta bursting rate, specifically at F3/P3 during encoding and F3 during delay, and HFA bursting duration, specifically F3 during encoding and P3 during delay, were most strongly associated with overall participant accuracy. Alpha findings were less consistent, as the bursting rate during the delay was negatively associated with accuracy at F3 and yet positively associated with accuracy at P3. This collectively suggests that optimal WM accuracy may involve beta in broadly facilitating cognitive control, HFA in encoding sensory content in a frontal-to-parietal feedback process, and alpha in inhibiting parietal and dis-inhibiting frontal cortex. Follow-up analyses of alpha rate, beta rate, and HFA duration examined the relationship to task accuracy across the scalp montage. Here, the beta burst rate findings localized to posterior regions during encoding and anterior regions during the delay, while no differences across the scalp montage for alpha and HFA. The HFA-ABC findings localized to posterior regions during both encoding and delay periods. This builds upon single-electrode findings to more clearly highlights the role of frontal beta bursting in successful WM. Certainly, this frontal beta bursting rate finding is highly consistent with prior WM research on the role of beta in cognitive control (Liljefors et al., 2023; Lundqvist et al., 2016), while the posterior HFA-ABC finding may relate to aspects of sensory encoding given prior parietal gamma findings(Murray et al., 2017; Roux et al., 2012). Interestingly, the relationship between HFA burst duration and task accuracy was strongest during low-medium (versus high) WM load, although no differences in other bands were found across WM load or WM demands. This indicates that the strength of this association does not increase with higher WM demands, and provides some evidence to suggest the relationship is specific to lower WM load.

Taken together, both on a trial-by-trial level and on the individual performance level, reduced alpha/beta-HFA dynamics were indicative of impaired performance. Although this is the first study to our knowledge that has shown incorrect versus correct trial bursting patterns in humans, it is consistent with prior non-human primate findings (Rassi et al.), as well as findings in adults showing that lower beta activity during the delay (compared to baseline) (Paulo et al., 2023) and across the whole trial (Pavlov & Kotchoubey, 2020) was associated with better WM performance. This suggests that the switching between these high and low oscillatory states during differing WM demands relates to task accuracy, and that less efficient switching between these states may be an underlying cause of WM deficits.

Our performance-related findings have critical implications for potential treatment targets of WM in clinical populations, such as schizophrenia, depression, and attention deficit hyperactivity disorder (ADHD). WM deficits are an established transdiagnostic component of all neuropsychiatric disorders (Willcutt, 2008) and WM is specifically identified as a risk factor for transdiagnostic psychopathology (Huang-Pollock et al., 2017). Despite being one of the strongest predictors of poor clinical and functional outcomes, there remains a dearth of available treatments for WM deficits (Diamond, 2013; Gardiner & Iarocci, 2018; Lee et al., 2013; Miller & Hinshaw, 2010; Rinsky & Hinshaw, 2011). Alpha/beta and gamma/HFA bursting dynamics may reflect promising biomarkers of such neuropsychiatric disorders that can be targeted and objectively measured. Targeting dysfunctional neural dynamics is now possible with currently available brain modulation paradigms such as transcranial magnetic stimulation (rTMS) and transcranial direct current stimulation (tDCS) (Widge & Miller, 2019). Meta-analytic findings have shown rTMS to the left dlPFC can improve WM in healthy adults and specific clinical samples, yet there remains a tremendous range in outcomes due to the non-specific targeting in traditional or clinical rTMS protocols (Brunoni & Vanderhasselt, 2014; Martin et al., 2016; Martin et al., 2017). However, it is now possible to send rTMS pulses via closed-loop protocols synced to real-time oscillatory dynamics and/or via task-locked protocols to specific pulses during specific neurocognitive demands (e.g., during encoding versus delay; (Widge & Miller, 2019). Current results indicate that overactive alpha/beta and underactive HFA during WM encoding or underactive alpha/beta and overactive HFA during WM delay is associated with WM errors, and therefore may be promising targets for closed loop or task-synced neuromodulation protocols.

Finally, a note on the limitations of the current study. Data utilized in this manuscript were made publicly available by the original authors (Pavlov & Kotchoubey, 2020) and utilized by our group to test current hypotheses. Given the 19-channel scalp EEG montage, source localization techniques were determined to be an inappropriate approach. Unlike more fine-tuned approaches with intracranial recordings, these findings are hampered by the poor spatial resolution and high degree of noise in human EEG recordings. At the same time, use of human EEG is critical to translating these scientific findings into actionable biomarkers in the clinical setting. Further, substantial line noise at 50 Hz was treated with a notch filter after other EEGLAB filtering approaches were unsuccessful, which may have interfered with accurate measuring of burst features in the higher frequency ranges (and why we restricted analyses to 55-80 Hz). Also note that in the absence of band-limited oscillatory activity in the gamma band, we here labelled activity in that range as HFA — comparing current HFA findings to prior gamma findings has inherent limitations given the nonoverlapping features of these dynamics. While correct/incorrect patterns were informative, data was obtained on a healthy young adult sample without psychiatric or neurological disorders. Follow-up work with clinical populations is needed in the process of translating findings to novel neurobiological treatment targets for WM deficits. As current results are purely correlational and causation cannot be implied, future studies should implement neuromodulatory techniques to probe a possible causal relationship. Additionally, while we examined HFA and relate our findings to prior gamma-band findings, we cannot distinguish underlying mechanisms, and it remains unclear whether HFA findings described here reflect interneuron-mediated mechanisms (as has been proposed by Miller and colleagues to describe narrow-band 30-100 Hz gamma bursting (Miller et al., 2018)) or alternatively, calcium/spike-dependent mechanisms. Due to our use of scalp EEG and the known signal to noise ratio, it is unclear whether the lack of bandlimited gamma activity reflects a lack of resolution of human EEG or differing neural phenomenon than that posted by Miller et al. Future work could build upon current findings to examine these dynamics with approaches that have a better signal to noise ratio (e.g., electrocorticography). Finally, future work should examine intra-individual relationships between trial level accuracy and bursting dynamics to more precisely identify person-specific markers of WM errors.

## Author Contributions

Brian Kavanaugh: Conceptualization, formal analysis, writing - original; Megan Vigne: Conceptualization, data curation, writing – review & editing; Christopher: Investigation, writing – review & editing; Ryan Thorpe: Conceptualization, writing – review & editing; Noah Vaughan: Formal analysis, writing – review & editing; W. Luke Acuff: Data curation, writing – review & editing; Eric Tirrell: Data curation, writing – review & editing; Saskia Haegens: Writing – review & editing, supervision; Linda Carpenter: Conceptualization, writing – review & editing, supervision; Stephanie Jones: Conceptualization, writing - original, supervision

## Data and Code Availability

We would like to thank Dr. Yuri Pavlov, PhD for making his EEG/WM dataset publicly available (https://nemar.org/dataexplorer/detail?dataset_id=ds003655) (Pavlov, 2020, 2021). All work described in this manuscript is based on Dr. Pavlov’s dataset. We are grateful for his emphasis on facilitating open science.

---

**Supplemental Table 1.** Summary of alpha, beta, and HFA findings at F3 and P3 during working memory demands.

| F3 |  |  |  |  |  |  |  |  |  |  |  |  |
| --- | --- | --- | --- | --- | --- | --- | --- | --- | --- | --- | --- | --- |
|  | Alpha |  |  |  | Beta |  |  |  | HFA |  |  |  |
|  | Rate | Power | Duration | F-Span | Rate | Power | Duration | F-Span | Rate | Power | Duration | F-Span |
| Changes with TFR | ✓ | ✓ | ✓ | ✓ | ✓ | ✓ |  | ✓ | ✓ | ✓ |  |  |
| Changes with Accuracy | ✓ |  |  |  | ✓ |  |  |  |  |  |  |  |
| Correlated with HFA or Alpha/Beta |  |  |  |  | ✓+ |  |  |  | ✓+ |  |  |  |
| P3 |  |  |  |  |  |  |  |  |  |  |  |  |
|  | Alpha |  |  |  | Beta |  |  |  | HFA |  |  |  |
|  | Rate | Power | Duration | F-Span | Rate | Power | Duration | F-Span | Rate | Power | Duration | F-Span |
| Changes with TFR | ✓ | ✓ | ✓ |  | ✓ | ✓ | ✓ |  | ✓ | ✓ |  |  |
| Changes with Accuracy | ✓ |  |  |  | ✓ | ✓ |  |  | ✓ |  |  |  |
| Correlated with HFA or Alpha/Beta | ✓- | ✓- | ✓- |  | ✓- | ✓- |  |  | ✓- |  |  |  |

Note. Checkmark denotes positive finding in the given band and burst feature. Gray coloring reflects changes associated with TFR. Green coloring reflects changes associated with task accuracy. Blue (positive association) and orange (negative association) coloring reflects association with the opposing band (i.e., alpha/beta correlates with HFA or HFA correlates with alpha/beta). TFR = Time frequency responses. HFA = High frequency activity (55-80 Hz).

**Supplemental Figure 1.**
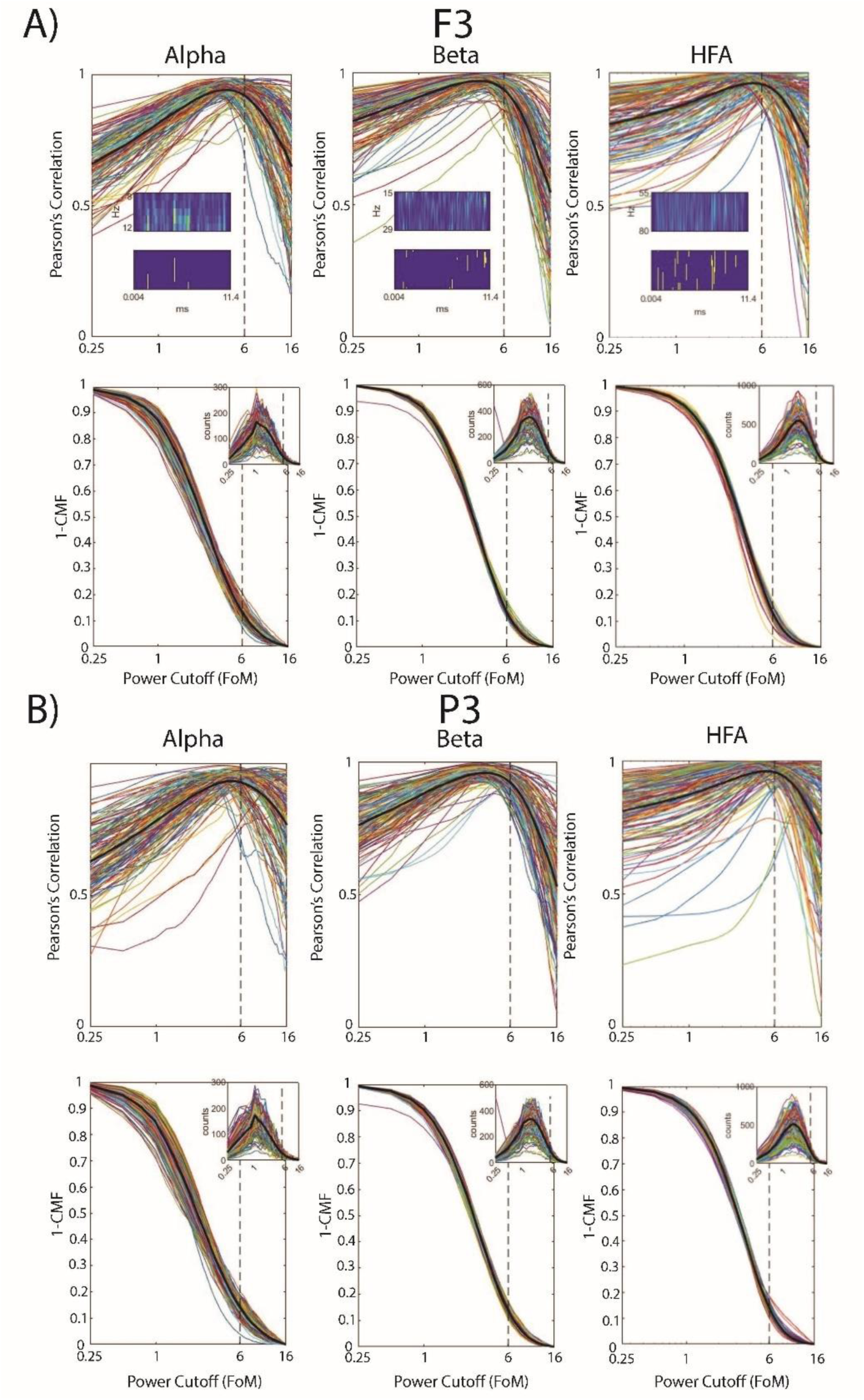
Bursts defined by a 6 factors-of-the-median (FOM) threshold show consistently high correlation with average spectral power within the band-of-interest (BOI) at F3 (A) and P3 (B). Across-trial Pearson’s correlation between time-averaged power within the BOI and percent area (i.e., percentage of pixels in the spectrogram) above the cutoff as a function of FOM threshold for each participant (colored traces) alongside the mean (black trace). Inset shows a representative spectrogram (upper) thresholded at 6 FOM (lower). Proportion of total possible bursts above the cutoff (1-CDF) as a function of FOM threshold for each participant (colored traces) alongside the aggregate (black trace). Inset shows the burst count histogram from which the cumulative distribution function (CDF) was calculated.

**Supplemental Figure 2.**
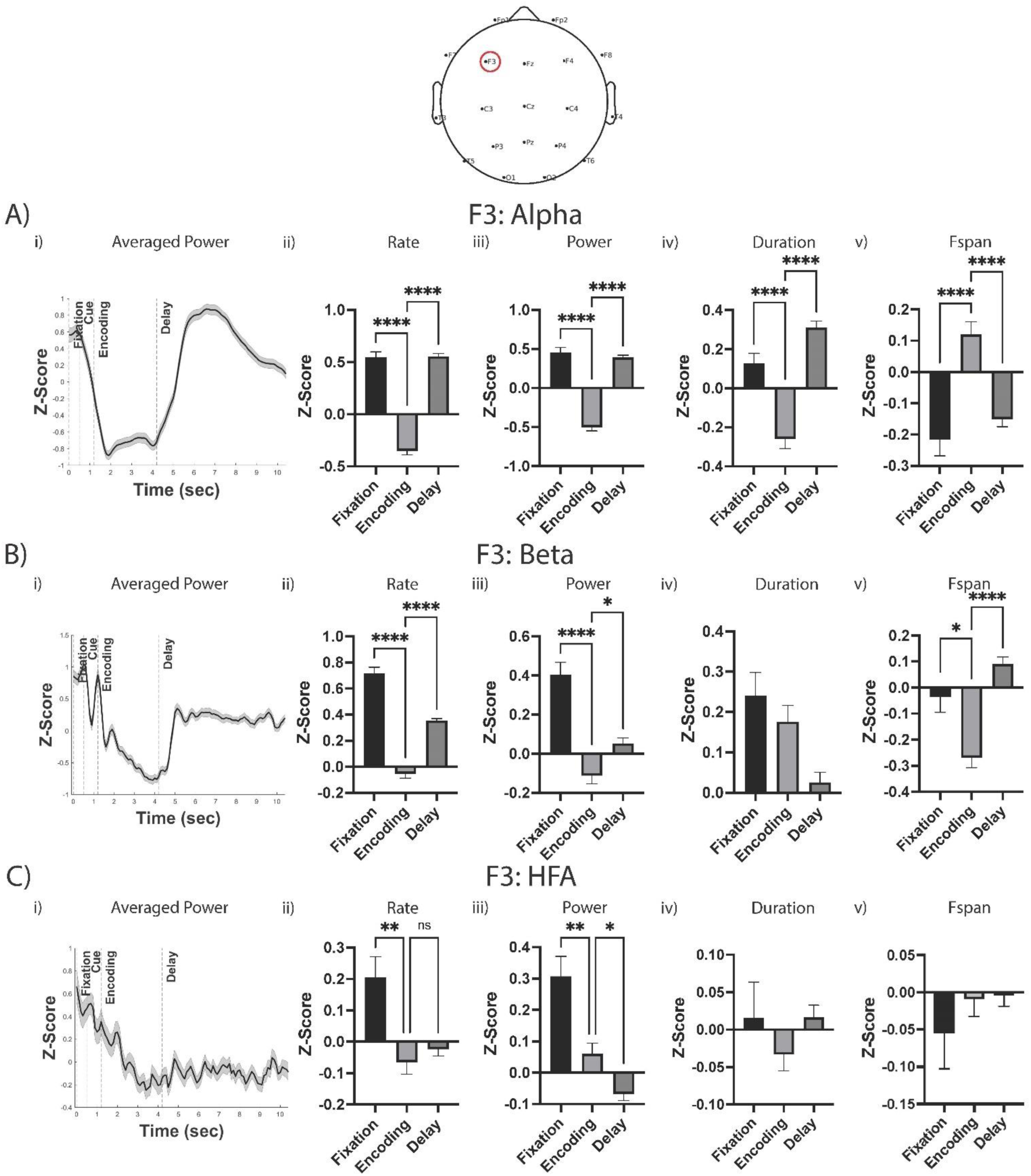
Burst power, duration, and frequency span, not just burst rate, change across WM stages in a pattern reflective of averaged power at F3. i) Averaged power time series across trial. ii-v) Oscillatory bursting patterns across fixation, encoding, and delay stages for burst rate (ii), burst power (iii), burst duration (iv), and burst frequency span (v) within alpha (A), beta (B), and HFA (C) bands. Asterisks denote statistical significance. Pairwise comparisons only displayed when overall model was statistically significant after Bonferroni correction.

**Supplemental Figure 3.**
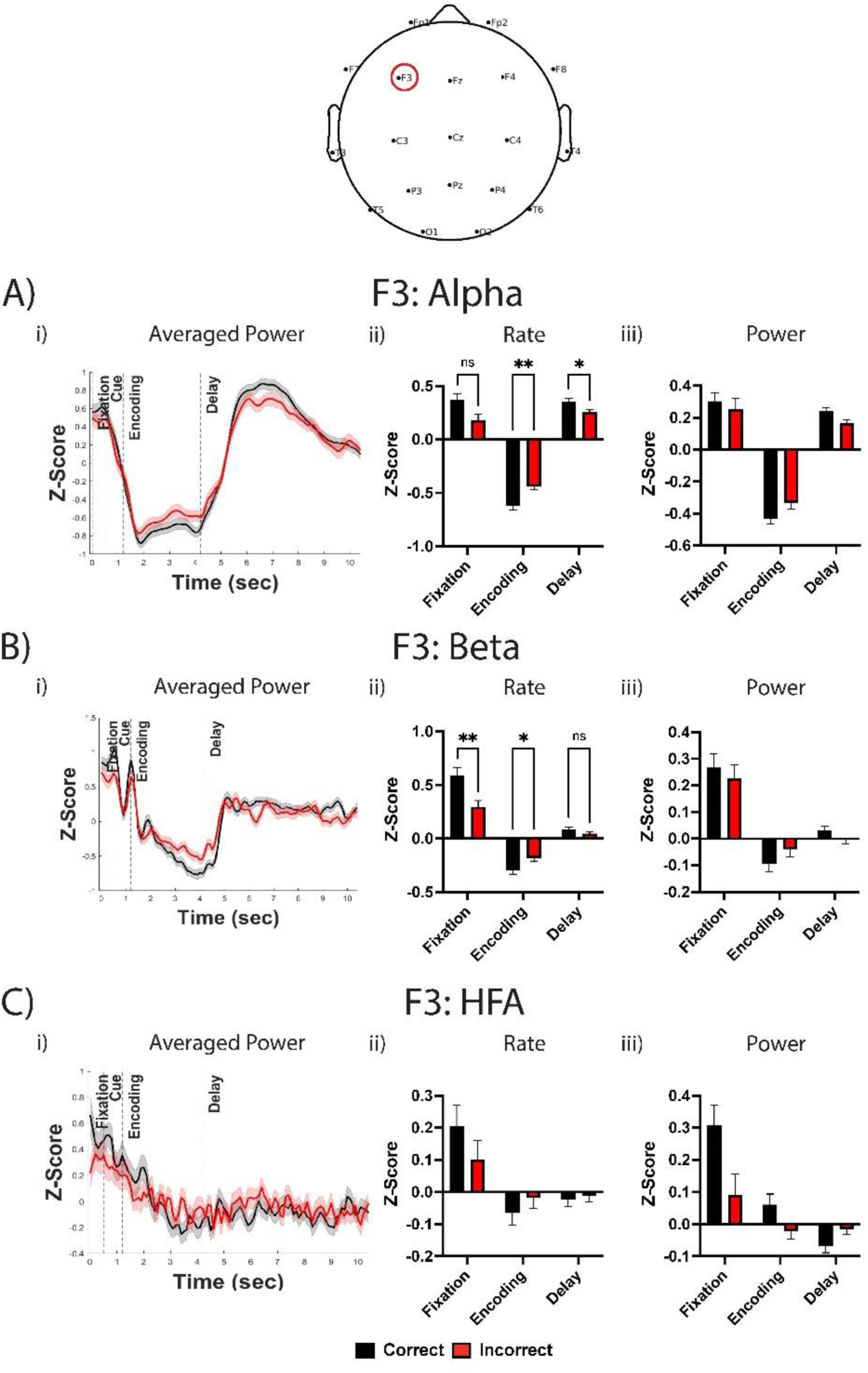
Incorrect responses were associated with diminished frontal bursting patterns across alpha, beta, and HFA bands. i) Averaged power time series across trial for correct and incorrect trials. ii-v) Time x accuracy analyses examining differences between electrodes in oscillatory bursting patterns across fixation, encoding, and delay stages for burst rate (ii) and burst power (iii) within alpha (A), beta (B), and HFA (C) bands. Asterisks denote statistical significance. Pairwise comparisons only displayed when overall model was statistically significant after Bonferroni correction.

**Supplemental Figure 4.**
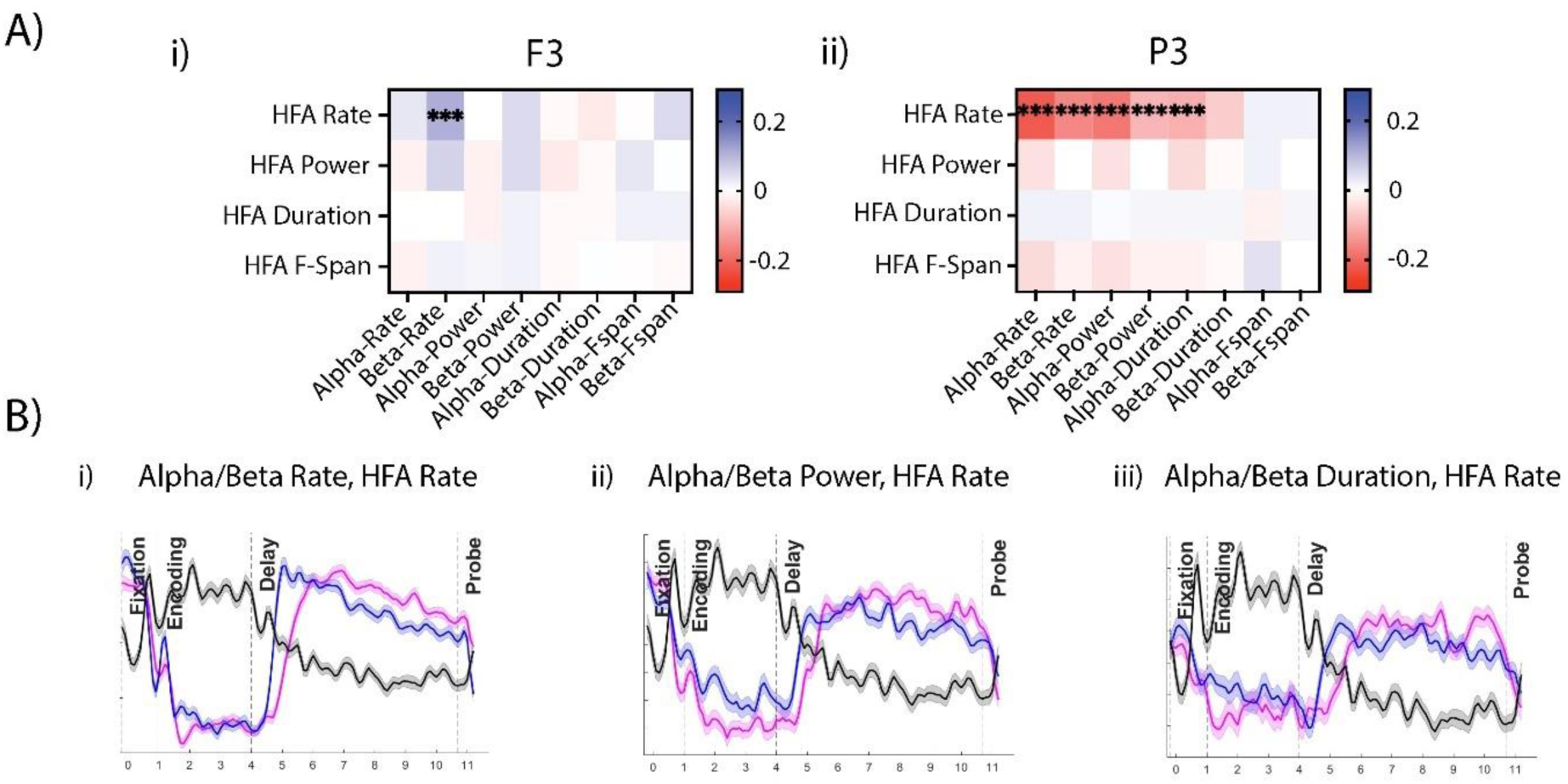
**A)** Correlation between HFA bursting and alpha/beta bursting at F3 (i) and P3 (ii). **B)** Blue = beta, red = alpha, black = HFA. Time x band analysis examining bursting patterns across time within alpha, beta, and HFA bands at P3 for HFA rate versus alpha/beta rate (i), alpha/beta power (ii), and alpha/beta duration (iii).

## REFERENCES

Brunoni, A. R., & Vanderhasselt, M. A. (2014). Working memory improvement with non-invasive brain stimulation of the dorsolateral prefrontal cortex: a systematic review and meta-analysis. Brain Cogn, 86, 1–9. 10.1016/j.bandc.2014.01.008

Buschman, T. J., & Miller, E. K. (2022). Working Memory Is Complex and Dynamic, Like Your Thoughts. J Cogn Neurosci, 35(1), 17–23. 10.1162/jocn_a_01940

Crowe, D. A., Goodwin, S. J., Blackman, R. K., Sakellaridi, S., Sponheim, S. R., MacDonald, A. W., 3rd, & Chafee, M. V. (2013). Prefrontal neurons transmit signals to parietal neurons that reflect executive control of cognition. Nat Neurosci, 16(10), 1484–1491. 10.1038/nn.3509

Diamond, A. (2013). Executive functions. Annu Rev Psychol, 64, 135–168. 10.1146/annurev-psych-113011-143750

ElShafei, H. A., Zhou, Y. J., & Haegens, S. (2022). Shaping Information Processing: The Role of Oscillatory Dynamics in a Working Memory Task. eNeuro, 9(5). 10.1523/ENEURO.0489-21.2022

Enz, N., Ruddy, K. L., Rueda-Delgado, L. M., & Whelan, R. (2021). Volume of beta-Bursts, But Not Their Rate, Predicts Successful Response Inhibition. J Neurosci, 41(23), 5069–5079. 10.1523/JNEUROSCI.2231-20.2021

Esterman, M., Chiu, Y., Tamber-Rosenau, B. J., & Yantis, S. (2009). Decoding cognitive control in human parietal cortex. PNAS, 106(42), 17974–17979.

Gardiner, E., & Iarocci, G. (2018). Everyday executive function predicts adaptive and internalizing behavior among children with and without autism spectrum disorder. Autism Res, 11(2), 284–295. 10.1002/aur.1877

Goodwin, S. J., Blackman, R. K., Sakellaridi, S., & Chafee, M. V. (2012). Executive control over cognition: stronger and earlier rule-based modulation of spatial category signals in prefrontal cortex relative to parietal cortex. J Neurosci, 32(10), 3499–3515. 10.1523/JNEUROSCI.3585-11.2012

Huang-Pollock, C., Shapiro, Z., Galloway-Long, H., & Weigard, A. (2017). Is Poor Working Memory a Transdiagnostic Risk Factor for Psychopathology? J Abnorm Child Psychol, 45(8), 1477–1490. 10.1007/s10802-016-0219-8

Iemi, L., Gwilliams, L., Samaha, J., Auksztulewicz, R., Cycowicz, Y. M., King, J. R., Nikulin, V. V., Thesen, T., Doyle, W., Devinsky, O., Schroeder, C. E., Melloni, L., & Haegens, S. (2022). Ongoing neural oscillations influence behavior and sensory representations by suppressing neuronal excitability. Neuroimage, 247, 118746. 10.1016/j.neuroimage.2021.118746

Jones, S. R. (2016). When brain rhythms aren’t ‘rhythmic’: implication for their mechanisms and meaning. Curr Opin Neurobiol, 40, 72–80. 10.1016/j.conb.2016.06.010

Kavanaugh, B. C., Fukuda, A. M., Gemelli, Z. T., Thorpe, R., Tirrell, E., Vigne, M., Jones, S. R., & Carpenter, L. L. (2023). Pre-treatment frontal beta events are associated with executive dysfunction improvement after repetitive transcranial magnetic stimulation for depression: A preliminary report. J Psychiatr Res, 168, 71–81. 10.1016/j.jpsychires.2023.10.024

Koenigs, M., Barbey, A. K., Postle, B. R., & Grafman, J. (2009). Superior parietal cortex is critical for the manipulation of information in working memory. J Neurosci, 29(47), 14980–14986. 10.1523/JNEUROSCI.3706-09.2009

Lara, A. H., & Wallis, J. D. (2015). The Role of Prefrontal Cortex in Working Memory: A Mini Review. Front Syst Neurosci, 9, 173. 10.3389/fnsys.2015.00173

Lee, R. S., Hermens, D. F., Redoblado-Hodge, M. A., Naismith, S. L., Porter, M. A., Kaur, M., White, D., Scott, E. M., & Hickie, I. B. (2013). Neuropsychological and socio-occupational functioning in young psychiatric outpatients: a longitudinal investigation. PLoS One, 8(3), e58176. 10.1371/journal.pone.0058176

Levitt, J., Edhi, M. M., Thorpe, R. V., Leung, J. W., Michishita, M., Koyama, S., Yoshikawa, S., Scarfo, K. A., Carayannopoulos, A. G., Gu, W., Srivastava, K. H., Clark, B. A., Esteller, R., Borton, D. A., Jones, S. R., & Saab, C. Y. (2020). Pain phenotypes classified by machine learning using electroencephalography features. Neuroimage, 223, 117256. 10.1016/j.neuroimage.2020.117256

Liljefors, J., Almeida, R., Rane, G., Lundström, J. N., Herman, P., & Lundqvist, M. (2023). 10.1101/2023.11.17.566386

Lundqvist, M., Herman, P., Warden, M. R., Brincat, S. L., & Miller, E. K. (2018). Gamma and beta bursts during working memory readout suggest roles in its volitional control. Nat Commun, 9(1), 394. 10.1038/s41467-017-02791-8

Lundqvist, M., Miller, E. K., Nordmark, J., Liljefors, J., & Herman, P. (2024). Beta: bursts of cognition. Trends Cogn Sci. 10.1016/j.tics.2024.03.010

Lundqvist, M., Rose, J., Herman, P., Brincat, S. L., Buschman, T. J., & Miller, E. K. (2016). Gamma and Beta Bursts Underlie Working Memory. Neuron, 90(1), 152–164. 10.1016/j.neuron.2016.02.028

Martin, D. M., McClintock, S. M., Forster, J., & Loo, C. K. (2016). Does Therapeutic Repetitive Transcranial Magnetic Stimulation Cause Cognitive Enhancing Effects in Patients with Neuropsychiatric Conditions? A Systematic Review and Meta-Analysis of Randomised Controlled Trials. Neuropsychol Rev, 26(3), 295–309. 10.1007/s11065-016-9325-1

Martin, D. M., McClintock, S. M., Forster, J. J., Lo, T. Y., & Loo, C. K. (2017). Cognitive enhancing effects of rTMS administered to the prefrontal cortex in patients with depression: A systematic review and meta-analysis of individual task effects. Depress Anxiety, 34(11), 1029–1039. 10.1002/da.22658

McKeon, S. D., Calabro, F., Thorpe, R. V., de la Fuente, A., Foran, W., Parr, A. C., Jones, S. R., & Luna, B. (2023). Age-related differences in transient gamma band activity during working memory maintenance through adolescence. Neuroimage, 274, 120112. 10.1016/j.neuroimage.2023.120112

Miller, E. K., Lundqvist, M., & Bastos, A. M. (2018). Working Memory 2.0. Neuron, 100(2), 463–475. 10.1016/j.neuron.2018.09.023

Miller, M., & Hinshaw, S. P. (2010). Does childhood executive function predict adolescent functional outcomes in girls with ADHD? J Abnorm Child Psychol, 38(3), 315–326. 10.1007/s10802-009-9369-2

Morris, A. T., Temereanca, S., Zandvakili, A., Thorpe, R., Sliva, D. D., Greenberg, B. D., Carpenter, L. L., Philip, N. S., & Jones, S. R. (2023). Fronto-central resting-state 15-29 Hz transient beta events change with therapeutic transcranial magnetic stimulation for posttraumatic stress disorder and major depressive disorder. Sci Rep, 13(1), 6366. 10.1038/s41598-023-32801-3

Murray, J. D., Jaramillo, J., & Wang, X. J. (2017). Working Memory and Decision-Making in a Frontoparietal Circuit Model. J Neurosci, 37(50), 12167–12186. 10.1523/JNEUROSCI.0343-17.2017

Niendam, T. A., Laird, A. R., Ray, K. L., Dean, Y. M., Glahn, D. C., & Carter, C. S. (2012). Meta-analytic evidence for a superordinate cognitive control network subserving diverse executive functions. Cogn Affect Behav Neurosci, 12(2), 241–268. 10.3758/s13415-011-0083-5

Paulo, D. L., Qian, H., Subramanian, D., Johnson, G. W., Zhao, Z., Hett, K., Kang, H., Chris Kao, C., Roy, N., Summers, J. E., Claassen, D. O., Dhima, K., & Bick, S. K. (2023). Corticostriatal beta oscillation changes associated with cognitive function in Parkinson’s disease. Brain, 146(9), 3662–3675. 10.1093/brain/awad206

Pavlov, Y. G., & Kotchoubey, B. (2020). The electrophysiological underpinnings of variation in verbal working memory capacity. Sci Rep, 10(1), 16090. 10.1038/s41598-020-72940-5

Pavlov, Y. G., & Kotchoubey, B. (2021). Temporally distinct oscillatory codes of retention and manipulation of verbal working memory. Eur J Neurosci, 54(7), 6497–6511. 10.1111/ejn.15457

Pavlov, Y. G., & Kotchoubey, B. (2022). Oscillatory brain activity and maintenance of verbal and visual working memory: A systematic review. Psychophysiology, 59(5), e13735. 10.1111/psyp.13735

Rassi, E., Zhang, Y., Mendoza, G., Mendez, J. C., Merchant, H., & Haegens, S. (2023). Distinct beta frequencies reflect categorical decisions. Nat Commun, 14(1), 2923. 10.1038/s41467-023-38675-3

Rinsky, J. R., & Hinshaw, S. P. (2011). Linkages between childhood executive functioning and adolescent social functioning and psychopathology in girls with ADHD. Child Neuropsychol, 17(4), 368–390. 10.1080/09297049.2010.544649

Rodriguez-Larios, J., & Haegens, S. (2023). Genuine beta bursts in human working memory: controlling for the influence of lower-frequency rhythms. bioRxiv. 10.1101/2023.05.26.542448

Roux, F., Wibral, M., Mohr, H. M., Singer, W., & Uhlhaas, P. J. (2012). Gamma-band activity in human prefrontal cortex codes for the number of relevant items maintained in working memory. J Neurosci, 32(36), 12411–12420. 10.1523/JNEUROSCI.0421-12.2012

Sherman, M. A., Lee, S., Law, R., Haegens, S., Thorn, C. A., Hamalainen, M. S., Moore, C. I., & Jones, S. R. (2016). Neural mechanisms of transient neocortical beta rhythms: Converging evidence from humans, computational modeling, monkeys, and mice. Proc Natl Acad Sci U S A, 113(33), E4885–4894. 10.1073/pnas.1604135113

Shin, H., Law, R., Tsutsui, S., Moore, C. I., & Jones, S. R. (2017). The rate of transient beta frequency events predicts behavior across tasks and species. Elife, 6. 10.7554/eLife.29086

Wang, X. J. (2010). Neurophysiological and computational principles of cortical rhythms in cognition. Physiol Rev, 90(3), 1195–1268. 10.1152/physrev.00035.2008

Wessel, J. R. (2020). beta-Bursts Reveal the Trial-to-Trial Dynamics of Movement Initiation and Cancellation. J Neurosci, 40(2), 411–423. 10.1523/JNEUROSCI.1887-19.2019

Widge, A. S., & Miller, E. K. (2019). Targeting Cognition and Networks Through Neural Oscillations: Next-Generation Clinical Brain Stimulation. JAMA Psychiatry, 76(7), 671–672. 10.1001/jamapsychiatry.2019.0740

Willcutt, E., Sonuga-Barke, E., Nigg, J., & Sergeant, J. (2008). Recent developments in neuropsychological models of childhood psychiatric disorders. Biological Child Psychiatry, 24, 195–226.

Zhou, Y. J., Ramchandran, A., & Haegens, S. (2023). Alpha oscillations protect working memory against distracters in a modality-specific way. Neuroimage, 278, 120290. 10.1016/j.neuroimage.2023.120290

